## Supplemental for "Time-varying stimuli that prolong IKK activation promote nuclear remodeling and mechanistic switching of NF-κB dynamics"

**Other footnotes**

**Supplementary files**

**Supplementary File 1: STL file for Y-channel microfluidic device.** The included file can be 3D printed and used as a mold to produce the PDMS-based microfluidic device used in this study.

**Supplementary File 2: D2FC$^2$ model files.** The included model files are used for D2FC$^2$ and related simulations. See also the lab GitHub page for most up to date information (https://github.com/recleelab/D2FCSquared/)

**Supplementary movies**

**Supplementary movie 1: Time-lapse images of CRISPR-modified U2OS cells.** Fluorescent protein fusions at their endogenous loci are shown EGFP-NEMO (left) and mCherry-RelA (right). 60x images captured the response of two cells exposed to continuous to a saturating 1000 ng/mL of IL-1.

**Supplementary movie 2: Time-lapse images of dual-reporter cells exposed to a single pulse.** Dynamics of EGFP-NEMO (left) and mCherry-RelA (right) in response to a single 6-minute pulse of 10 ng/mL IL-1 in the microfluidic device. Cells display typical adaptive behavior within 100 minutes of stimulation.

**Supplementary movie 3: Time-lapse images of dual-reporter cells exposed to four 1.5-minute pulses.** Dynamics of EGFP-NEMO (left) and mCherry-RelA (right) in the microfluidic device exposed to 4x1.5-minute pulse of 10 ng/mL IL-1 with 5-minute gaps. Top cells displays prolonged EGFP-NEMO puncta and zero-order export kinetics, whereas the lower cell displays less-sustained EGFP-NEMO puncta and adapts within 180 minutes of stimulation.

**Supplementary figures**

**Supplementary Figure 1: Dynamic stimulation device and cell responses to a single pulse**

**A.** Schematic of the microfluidic chip as designed in the Autodesk Fusion 360 software highlighting the dimensions of the chip for the X, Y, and Z axis, units in mm. **B.** Image of the microfluidic chip with PDMS attached to a glass cover slide, coin for scale. Green food coloring was imaged to highlight the inlets, chamber, and outlets for image purposes only. **C.** Nuclear RelA export scores (see methods) for experimental trajectories of varying single pulse duration. **D.** Pairwise comparison of p-values for conditions in (C), student's t-test.

Figure S1

A

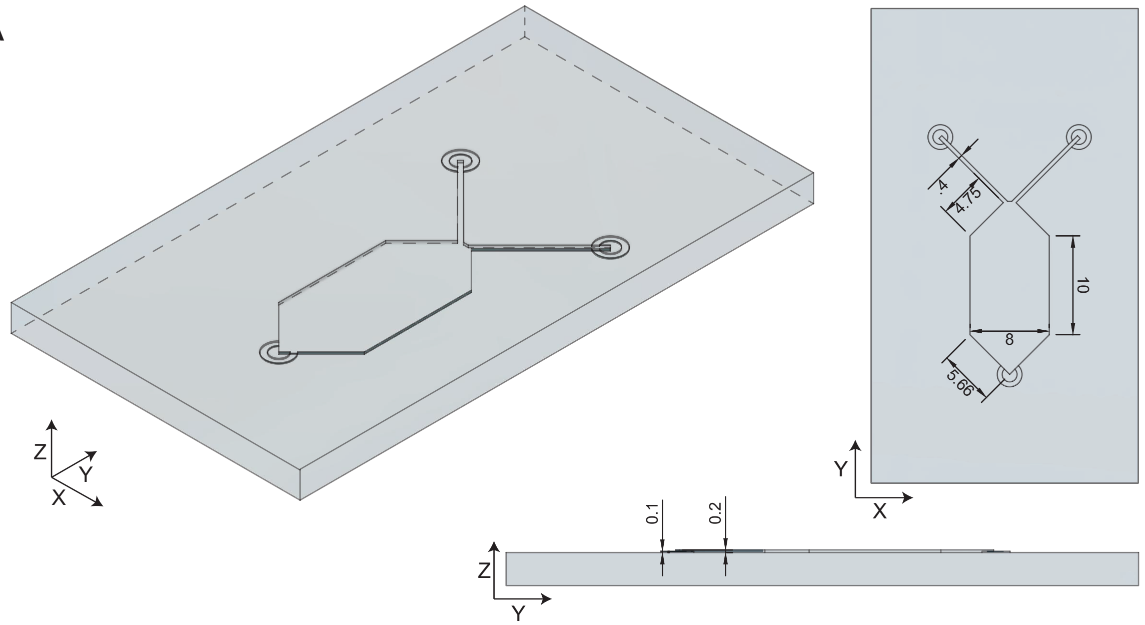


B

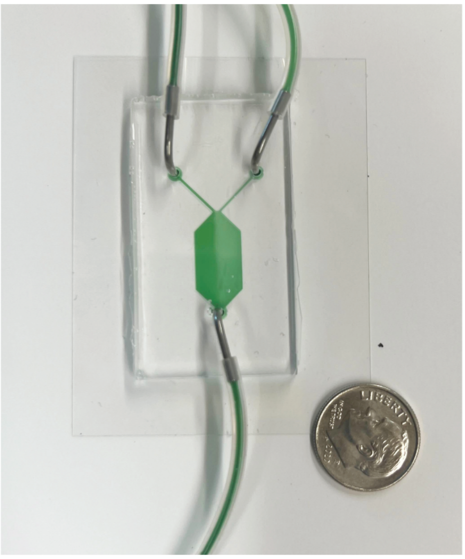

C

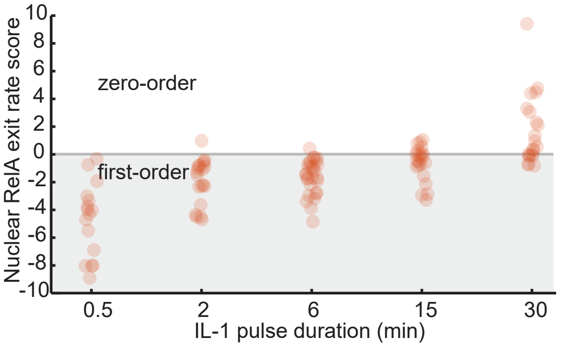


D

| p-val | 0.5 min | 2 min | 6 min | 15 min | 30 min |
|---|---|---|---|---|---|
| 0.5 min | 1 | 0.0002 | 3E-06 | 6E-07 | 1E-08 |
| 2 min | 0.0002 | 1 | 0.4 | 0.004 | 3E-06 |
| 6 min | 3E-06 | 0.4 | 1 | 0.007 | 3E-07 |
| 15 min | 6E-07 | 0.004 | 0.007 | 1 | 0.0005 |
| 30 min | 1E-08 | 3E-06 | 3E-07 | 0.0005 | 1 |

**Supplementary Figure 2: NF-kB responses to a dose-conserving cytokine pulse train are enhanced for two and three pulse stimuli.**

**A.** Boxplots of single-cell experimental time courses showing AUCs of NEMO spots (left) and Nuclear RelA (right). A 5-minute gap time between three 2-minute pulses of IL-1 at 10 ng/mL causes a significant enhancement of nuclear RelA. Increasing the gap time beyond 5-minutes has diminishing returns. **B.** Boxplots of single-cell experimental time courses as in panel A for two 3-minute pulses. Increasing the gap duration up to 15 minutes enhances the AUC of nuclear RelA. Boxplots show the median value and interquartile ranges. On average, 20 cells were imaged for each condition. P-values indicated for student's t-test.

Figure S2

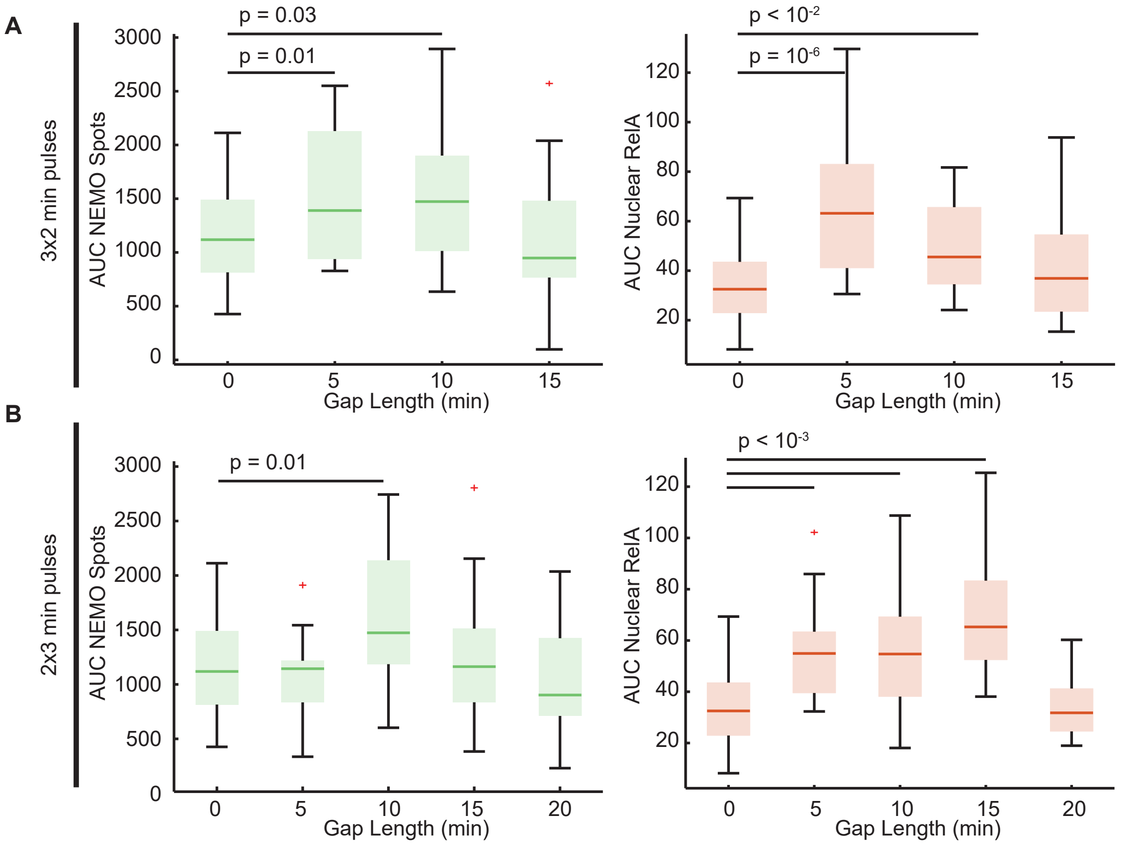

A
3x2 min pulses
AUC NEMO Spots
p = 0.03
p = 0.01
3000
2500
2000
1500
1000
500
0
0
5
10
15
Gap Length (min)
AUC Nuclear RelA
p < 10-2
p = 10-6
120
100
80
60
40
20
0
5
10
15
Gap Length (min)
B
2x3 min pulses
AUC NEMO Spots
p = 0.01
3000
2500
2000
1500
1000
500
0
0
5
10
15
20
Gap Length (min)
AUC Nuclear RelA
p < 10-3
120
100
80
60
40
20
0
5
10
15
20
Gap Length (min)

**Supplementary Figure 3: KYM-1 cells exposed to a pulse train of TNF enhances the NF-kB response**

**A.** Time lapse images of KYM-1 stably expressing mVen-RelA fusion exposed to TNF at 1 ng/mL as a single 6-minute pulse (top) or two 3-minute pulses with a 9-minute gap (bottom). **B.** Single cell trajectories of the intensity of nuclear RelA (fold change) for KYM-1 cells exposed to a continuous (left), single 6-minute (middle), and two 3-minute (right) pulses of TNF at 1 ng/mL. Orange bold lines represent mean and shaded region represents +/- 1 standard deviation of single cell trajectories. **C.** Box plots of the AUC of nuclear RelA (fold change) for varying the gap length of two 3-minute pulses (left) and two 2-minute pulses. Boxplots show the median value and interquartile ranges.

Figure S3

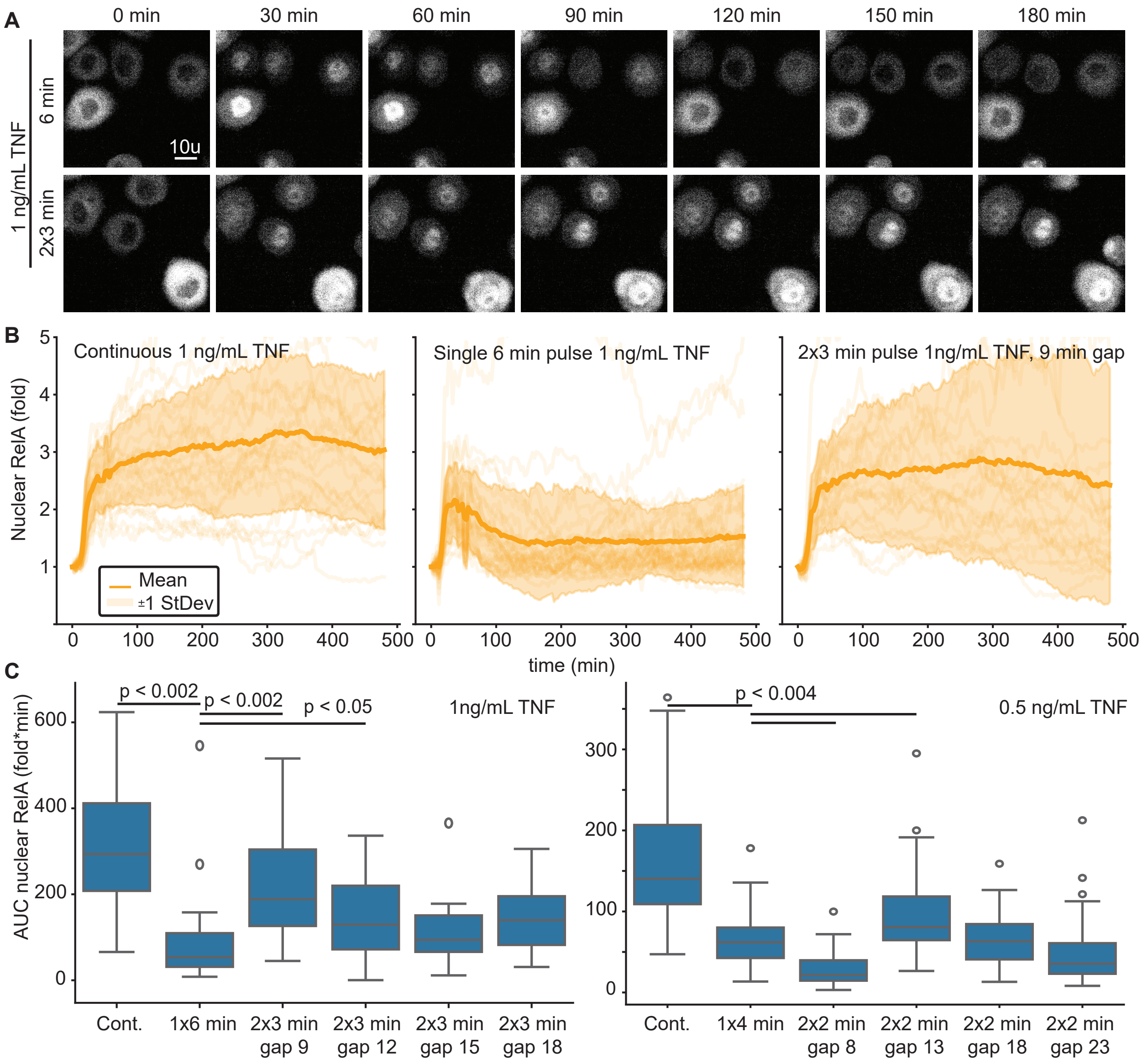
A
0 min
30 min
60 min
90 min
120 min
150 min
180 min
1 ng/mL TNF
6 min
2x3 min
10u
B
Continuous 1 ng/mL TNF
Single 6 min pulse 1 ng/mL TNF
2x3 min pulse 1ng/mL TNF, 9 min gap
Nuclear RelA (fold)
Mean
±1 StDev
time (min)
C
AUC nuclear RelA (fold*min)
p < 0.002
p < 0.002
p < 0.05
1ng/mL TNF
Cont.
1x6 min
2x3 min gap 9
2x3 min gap 12
2x3 min gap 15
2x3 min gap 18
p < 0.004
0.5 ng/mL TNF
Cont.
1x4 min
2x2 min gap 8
2x2 min gap 13
2x2 min gap 18
2x2 min gap 23

**Supplementary Figure 4: Dose-conserving pulse trains switch the export mechanism of nuclear RelA to a pseudo-zero-order process.**

**A.** Box plot of the maximum nuclear RelA (fold change) of single cell trajectories given 10 ng/ml IL-1 for single and multiple short pulses. Although significant for 2 of the 5 conditions, the trend of increased nuclear RelA is only very subtle and does not explain the enhanced AUC in response to a cytokine pulse train. **B.** Example trajectories highlighting the exit rate score, used to distinguish first-order and zero-order export kinetics. Experimental data are representative single cell trajectories (solid red line) and dashed lines represent models of best fit for zero and first order decay. Positive scores represent zero-order export kinetics, and negative scores represent first-order kinetics. See methods for details. **C.** Jitter plot of exit rate scores for individual cell trajectories of the single- and multi-pulse IL-1 stimulation patterns (top). Pairwise comparisons indicate p-value for student's t-test (bottom).

Figure S4

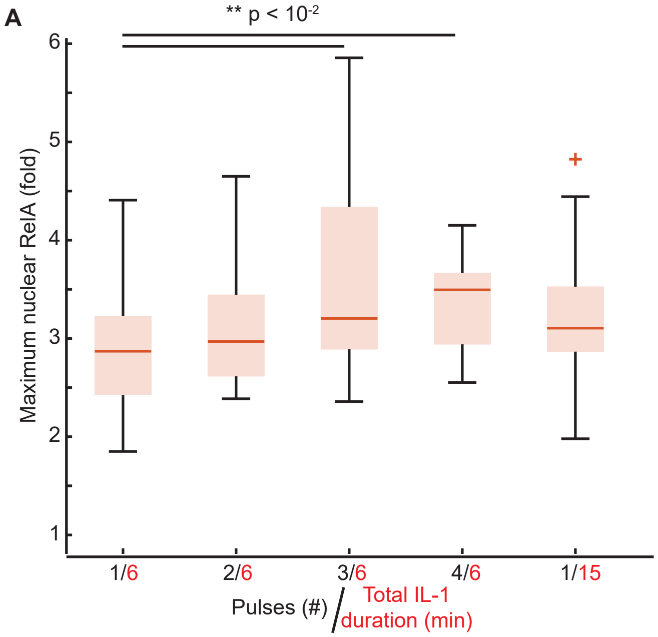


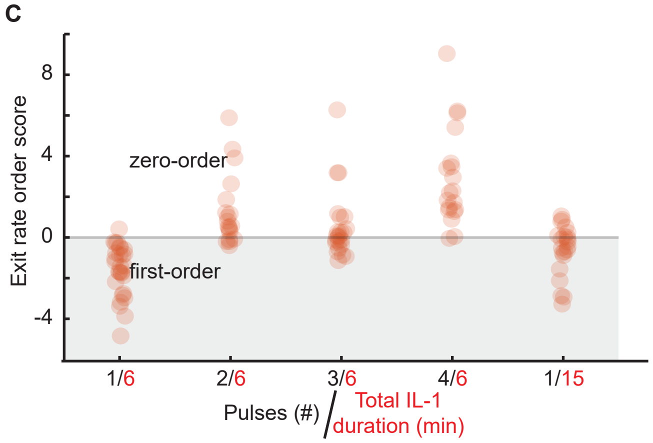


| | 1x6 min | 2x3 min | 3x2 min | 4x1.5 min | 1x15 min |
|---|---|---|---|---|---|
| 1x6 min | 1 | 4E - 08 | 3E-06 | 7E - 11 | 0.007 |
| 2x3 min | 4E-08 | 1 | 0.1 | 0.02 | 0.0002 |
| 3x2 min | 3E-06 | 0.1 | 1 | 0.0005 | 0.01 |
| 4x1.5 min | 7E-11 | 0.02 | 0.0005 | 1 | 4E - 07 |
| 1x15 min | 0.007 | 0.0002 | 0.01 | 4E - 07 | 1 |

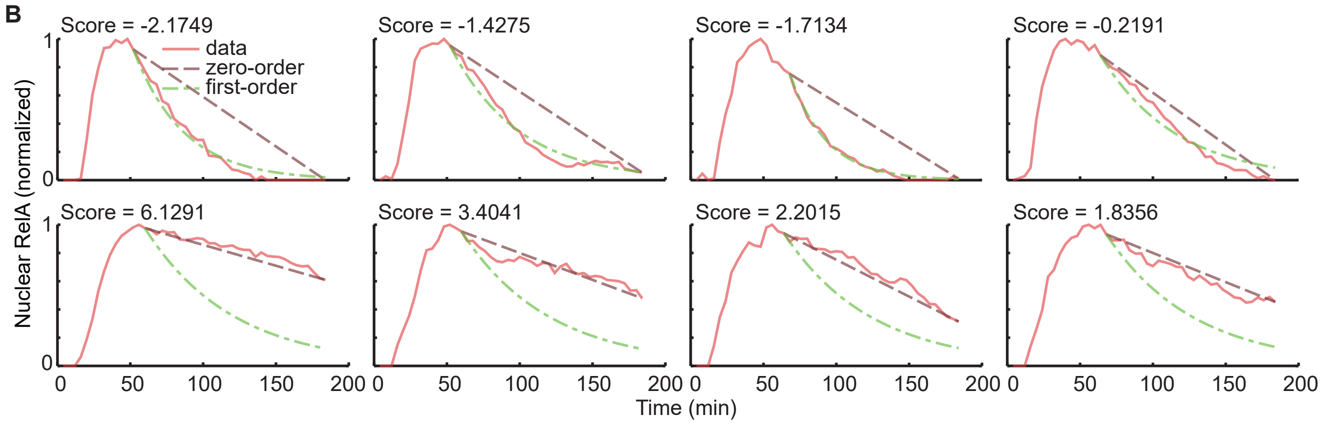

**Supplementary Figure 5: Original D2FC and optimized parameterizations do not recapitulate the emergent property.**

**A and B.** Box plots comparing the D2FC simulated results of the AUC nuclear RelA of the single (A) and multi-pulse (B) stimulation pattern. D2FC model is using original parameter set and D2FC optimized is the overall best set of parameters found from the particle swarm optimization. Statistically significant differences between models and experiment are indicated (*; $p<0.05$, student's t-test). The D2FC recapitulates responses to a single pulse, but not to pulse trains. Remarkably, the optimized D2FC fails to recapitulate most conditions. The relative failure of the optimized D2FC is because PSO using the emergent property (fits to averages from four selected conditions: control, 1x0.5-min, 4x1.5-min, 1x30-min) as an objective function effectively distributed the error across conditions, resulting in a model that marginally improves responses to a pulse train at the expense of even poorer fits to other conditions. **C.** Results of the D2FC, D2FC optimized, and $D2FC^2$ comparing simulated results (solid lines) with the averages of single-cell trajectories (circles) in each of three conditions. Simulation results for the D2FC optimized, and $D2FC^2$ are for lowest-error parameterizations following particle swarm optimization.

Figure S5

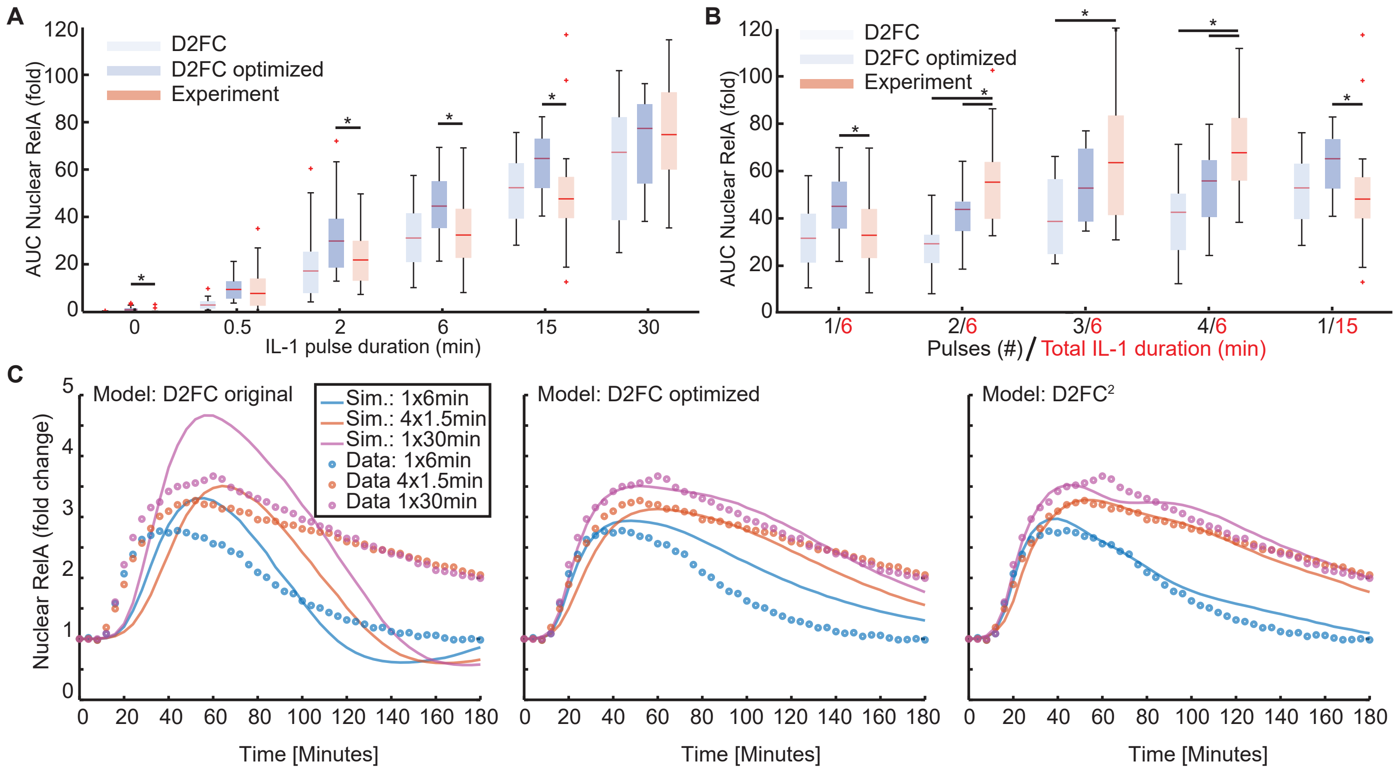
A
D2FC
D2FC optimized
Experiment
AUC Nuclear RelA (fold)
IL-1 pulse duration (min)
B
D2FC
D2FC optimized
Experiment
AUC Nuclear RelA (fold)
1/6
2/6
3/6
4/6
1/15
Pulses (#) / Total IL-1 duration (min)
C
Model: D2FC original
Sim.: 1x6min
Sim.: 4x1.5min
Sim.: 1x30min
Data: 1x6min
Data 4x1.5min
Data 1x30min
Nuclear RelA (fold change)
Time [Minutes]
Model: D2FC optimized
Time [Minutes]
Model: D2FC²
Time [Minutes]

**Supplementary Figure 6: Post-hoc analysis of particle swarm optimization results for the D2FC$^2$.**

**A.** Box plots of the parameters identified in 500 iterations of the D2FC$^2$ optimization using particle swarm. The y-axis is normalized relative to the parameter limits to account for differences in parameter orders. Importantly, for most parameters, there were solutions following particle swarm optimization that used the full range of values from the prior distribution suggesting that the parameter space has been effectively swarmed. **B.** Heat map of parameters for the top 10 models for the D2FC$^2$ normalized to the explored parameter space. Colorbar represents the normalized parameter value relative to the prior distribution (Supplementary Table 4). For most parameters there is variance between the models indicating that comparable results can be achieved via distinct parameter combinations.

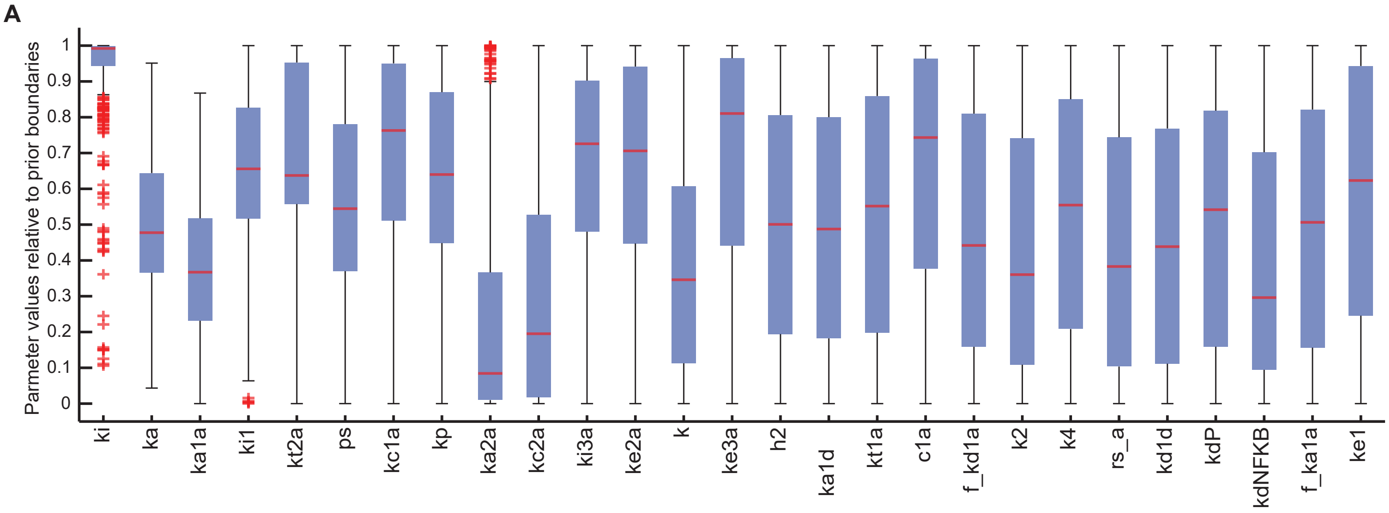
A
Parmeter values relative to prior boundaries
ki
ka
ka1a
ki1
kt2a
ps
kc1a
kp
ka2a
kc2a
ki3a
ke2a
k
ke3a
h2
ka1d
kt1a
c1a
f_kd1a
k2
k4
rs_a
kd1d
kdP
kdNFKB
f_ka1a
ke1

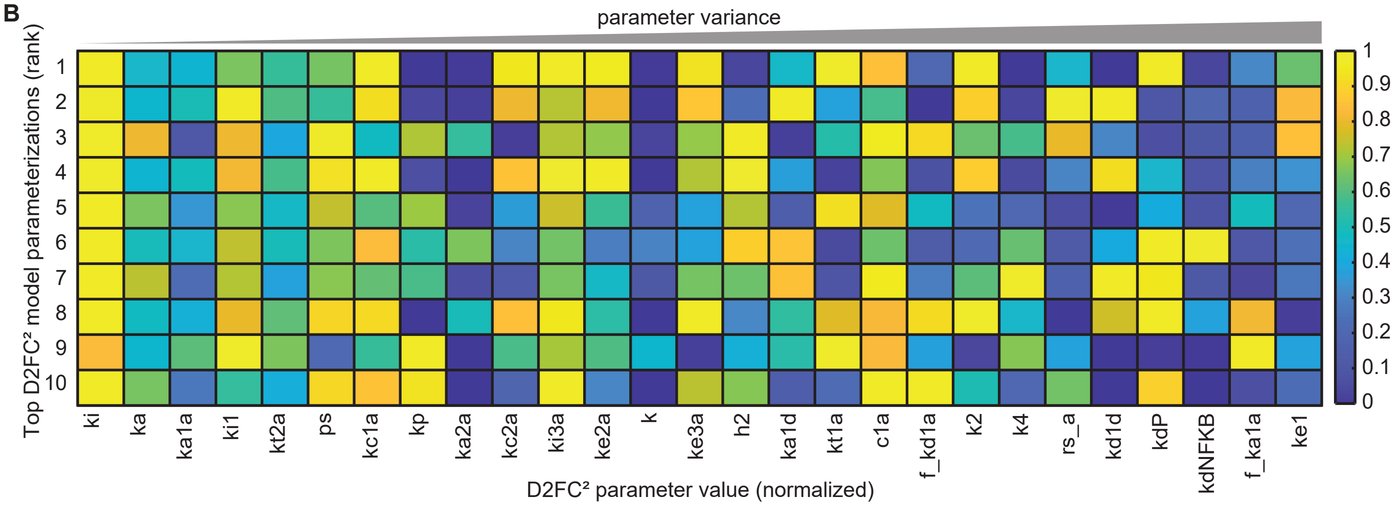
B
parameter variance
Top D2FC² model parameterizations (rank)
D2FC² parameter value (normalized)
ki
ka
ka1a
ki1
kt2a
ps
kc1a
kp
ka2a
kc2a
ki3a
ke2a
k
ke3a
h2
ka1d
kt1a
c1a
f_kd1a
k2
k4
rs_a
kd1d
kdP
kdNFKB
f_ka1a
ke1

**Supplementary Figure 7: Quality of fit to single cell trajectories for the top D2FC$^2$ model parameterizations.**

**A.** Elbow plot showing the fraction of non-acceptable fits as a function of the threshold for defining an acceptable single-cell trajectory. The elbow point, mathematically identified as the point furthest from a linear line connecting the first and last values (see Methods), was used to set a threshold between high- and poor-quality fits. A strict threshold of half the elbow point was applied to identify single cell simulations fits that are considered excellent.

**B.** Example simulated single cell trajectories highlighting the excellent, high, and poor single cell trajectories compared to the experimental data.

Figure S7

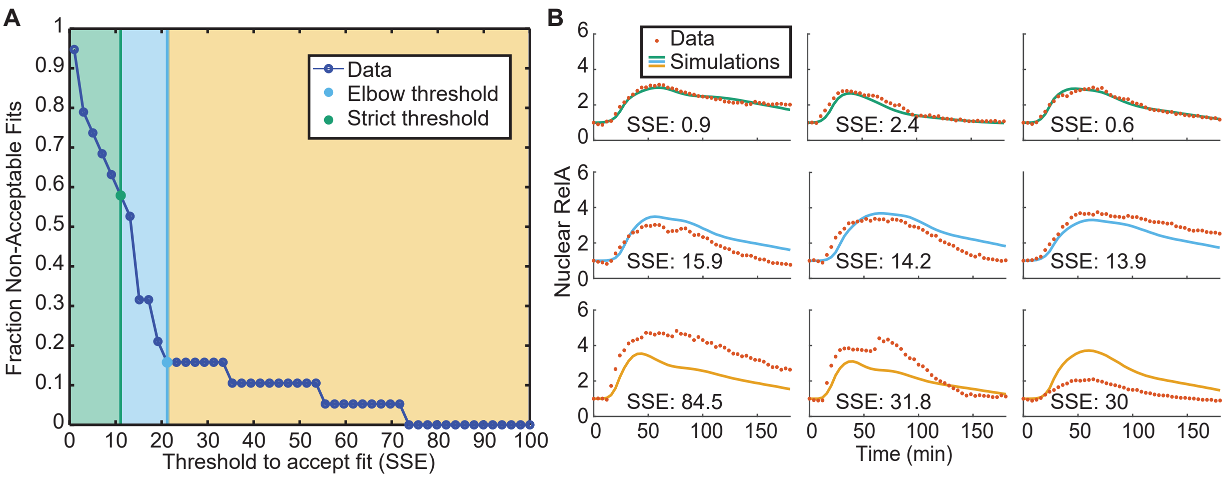

A
Fraction Non-Acceptable Fits
Threshold to accept fit (SSE)
Data
Elbow threshold
Strict threshold
B
Data
Simulations
Nuclear RelA
Time (min)
SSE: 0.9
SSE: 2.4
SSE: 0.6
SSE: 15.9
SSE: 14.2
SSE: 13.9
SSE: 84.5
SSE: 31.8
SSE: 30

**Supplementary Figure 8: 5-Azacytidine stimulation significantly reduces features of NF-kB responses.**

**A.** Box plots of the maximum nuclear RelA (fold change) for single cell trajectories exposed to 5-azacytidine for indicated duration before exposure to a single 6-minute pulse of 10 ng/ml IL-1. **B.** Box plots of the AUC for nuclear RelA (fold change) for single cell trajectories exposed to 5-azacytidine for indicated duration before exposure to a single 6-minute pulse of 10 ng/ml IL-1. P-values represent a student's t-test.

Figure S8

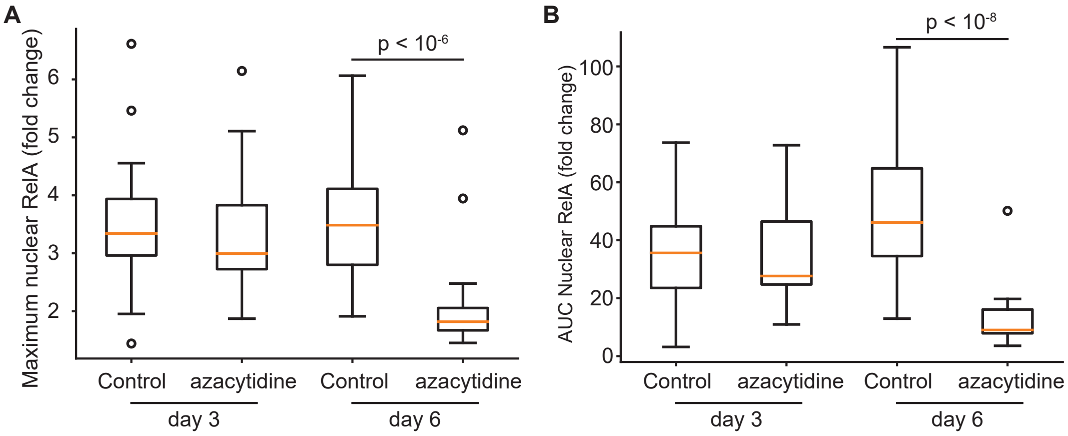

A
Maximum nuclear RelA (fold change)
6
5
4
3
2
p < 10-6
Control
azacytidine
Control
azacytidine
day 3
day 6
B
AUC Nuclear RelA (fold change)
100
80
60
40
20
0
p < 10-8
Control
azacytidine
Control
azacytidine
day 3
day 6

### Supplementary tables

**Supplementary Table 1:** Summarizes reactions for the D2FC² model where "C." and "N." prefixes denote species is in the cytoplasm and nucleus, respectively. The column labeled "name" represents the parameter influencing the reaction rate. The reported values are from the best D2FC² model. The rows highlighted in gray represent added model mechanisms that differentiate between D2FC and D2FC². Parameter kv (yellow; value = 3.3) accounts for subcellular compartment volumes between the cytoplasm and nucleus, applied to species traveling between the compartments. DCoop, NPio, NEnh1, NEnh2, and IKKSpots(t) are functions defined in supplementary table 2. Parameter ka (red) was used to optimize the original D2FC model to interface directly with NEMO spot numbers.

| Reaction | Name | Value | Rate |
|---|---|---|---|
| $C.IkBa + C.NFkB \rightarrow C.IkBaNFkB$ | ka1a | 5.854E-01 | $ka1a * [C.IkBa] * [C.NFkB]$ |
| $C.IkBaNFkB \rightarrow C.IkBa + C.NFkB$ | kd1a | 5.000E-02 | $kd1a * [C.IkBaNFkB]$ |
| $C.NFkB \rightarrow N.NFkB$ | ki1 | 5.403E-03 | $kv * ki1 * [C.NFkB]$ |
| $N.NFkB \rightarrow C.NFkB$ | ke1 | 9.657E-05 | $kv * ke1 * [N.NFkB]$ |
| $N.NFkB + N.IkBa \rightarrow N.IkBaNFkB$ | f_ka1a | 4.25E-02 | $f\_ka1a * ka1a * [N.NFkB] * [N.IkBa]$ |
| $N.IkBaNFkB \rightarrow N.NFkB + N.IkBa$ | f_kd1a | 2.625E-02 | $f\_kd1a * kd1a * [N.IkBaNFkB]$ |
| $N.NFkB \rightarrow N.NFkBDNA$ | ka1d | 7.070E-04 | $ka1d * DCoop * NPio * [N.NFkB]$ |
| $N.NFkBDNA \rightarrow N.NFkB$ | kd1d | 6.684E-05 | $kd1d * [N.NFkBDNA]$ |
| $N.IkBa + N.NFkBDNA \rightarrow N.IkBaNFkB$ | ka2a | 1.320E-02 | $ka2a * [N.IkBa] * [N.NFkBDNA]$ |
| $N.IkBa \rightarrow \emptyset$ | c4a | 5.000E-04 | $c4a * [N.IkBa]$ |
| $C.IkBa \rightarrow \emptyset$ | c4a | 5.000E-04 | $c4a * [C.IkBa]$ |
| $N.IkBaNFkB \rightarrow C.IkBaNFkB$ | ke2a | 8.353E-02 | $kv * ke2a * [N.IkBaNFkB]$ |
| $C.IkBa \rightarrow N.IkBa$ | ki3a | 6.638E-03 | $kv * ki3a * [C.IkBa]$ |
| $N.IkBa \rightarrow C.IkBa$ | ke3a | 2.457E-03 | $kv * ke3a * [N.IkBa]$ |
| $C.IkBaNFkB \rightarrow C.NFkB$ | c5a | 2.200E-05 | $c5a * [C.IkBaNFkB]$ |
| $C.IKK \rightarrow C.IKKi$ | ki | 1.156E-02 | $ki * [C.IKK]$ |
| $C.IKKi \rightarrow C.IKKn$ | kp | 6.464E-05 | $kp * [C.IKKi] * A20I$ |
| $C.IKKn \rightarrow C.IKK$ | ka | 2.409E-06 | $\mathrm{TR} * \mathrm{ka} * \mathrm{IKKSpots(t)} * [C.IKKn]$ |
| $\emptyset \rightarrow C.tIkBa$ | rs_a | 6.852E-08 | $rs\_a * (1 + NEnh1)$ |
| $C.tIkBa \rightarrow \emptyset$ | c3a | 3.000E-04 | $c3a * [C.tIkBa]$ |
| $\emptyset \rightarrow C.IkBa$ | c2a | 5.000E-01 | $c2a * C.tIkBa$ |
| $\emptyset \rightarrow C.tCompetitor$ | rs_a | 6.852E-08 | $rs\_a * c1a * NEnh2$ |
| $C.tCompetitor \rightarrow \emptyset$ | c6a | 4.290E-05 | $c6a * [C.tCompetitor]$ |
| $\emptyset \rightarrow C.Competitor$ | c2a | 5.000E-01 | $c2a * [C.tCompetitor]$ |
| $C.Competitor \rightarrow \emptyset$ | c4a | 5.000E-04 | $c4a * [C.Competitor]$ |
| $\emptyset \rightarrow C.tA20$ | c1 | 2.000E-07 | $c1 * NEen3$ |
| $C.tA20 \rightarrow \emptyset$ | c3 | 4.000E-04 | $c3 * [C.Competitor]$ |
| $\emptyset \rightarrow C.A20$ | c2 | 5.000E-01 | $c2 * [C.A20]$ |
| $C.A20 \rightarrow \emptyset$ | c4 | 4.500E-03 | $c4 * [C.A20]$ |
| $C.IkBa \rightarrow C.pIkBa$ | kc1a | 4.998E-01 | $kc1a * [C.IKK] * [C.IkBa]$ |
| $C.pIkBa \rightarrow \emptyset$ | kt1a | 5.796E-03 | $kt1a * [C.pIkBa]$ |
| $C.IkBaNFkB \rightarrow pIkBaNFkB$ | kc2a | 4.071E-01 | $kc2a * [C.IKK] * [C.IkBaNFkB]$ |
| $C.pIkBaNFkB \rightarrow C.NFkB$ | kt2a | 1.274E-04 | $kt2a * [C.pIkBaNFkB]$ |

**Supplementary Table 2:** Algebraic equations used in the ordinary differential equations as shown in Supplementary Table 1 and used as outputs to the model.

| Name | Equation | Notes |
|---|---|---|
| $DCoop$ | $\frac{[N.NFkB]^{h2}}{(kdNFkB)^{h2} + [N.NFkB]^{h2}}$ | Represents the cooperative effects of NFkB binding to DNA. |
| $NPio$ | $1 + ps * \frac{[N.NFkBDNA]^{h3}}{(kdP)^{h3} + [N.NFkBDNA]^{h3}}$ | Represents pioneering effect of NFKB can have on increasing the availed sites |
| $A20I$ | $\frac{kbA20}{kbA20 + C.A20}$ | Decrease rate of inactivated IKK converting to neutral IKK. |
| IKKSpots(t) | $IKK(t) = \sum_{i=1}^{4} a_i e^{\left[-\left(\frac{(x-b_i)}{c_i}\right)^2\right]}$ | Parameters $a_i$, $b_i$, and $c_i$ are fit to an experimental IKK spot profile. See supplementary information for complete set of parameters. |
| $NEnh1$ | $c1a * \frac{[N.NFkB]^h}{k^h + [N.NFkB]^h}$ | Increased rate of IkB transcription due to NFkB retained in nucleus. |
| $NEnh2$ | $c1a * \frac{\left(\frac{[N.NFkB]}{k}\right)^{h+1}}{1 + \left(\frac{[N.NFkB]}{k}\right)^{h+1} + \left(\frac{[Competitor]}{k4}\right)^{h+1}}$ | Increased rate of Competitor transcription where the competitor itself can reduce its own rate of transcription. |
| $NEen3$ | $c1 * \frac{\left(\frac{[N.NFkB]}{k}\right)^{h+1}}{1 + \left(\frac{[N.NFkB]}{k}\right)^{h+1} + \left(\frac{[Competitor]}{k2}\right)^{h+1}}$ | Increased rate of A20 transcription deepened upon NFkB nuclear localization and the rate can be reduced by competitor formation. |
| $N.NFkB_{Total}$ | $[N.NFkB] + [N.IkBaNFkB] + [N.NFkBDNA]$ | Total amount of NFkB in the nucleus including all forms which include free, bound to IkBa and bound to DNA |
| NFkB (Fold Change) | $\frac{[N.NFkB_{Total}]}{N.NFkB_{Total}(t=0)}$ | NFKB fold change that is directly comparable to the single cell data. |

**Supplementary Table 3:** Parameterizations of the top 10 D2FC$^2$ models found through particle swarm optimization.

| | Top 10 D2FC$^2$ parameterizations | | | | | | | | | |
|---|---|---|---|---|---|---|---|---|---|---|
| Name | 1 | 2 | 3 | 4 | 5 | 6 | 7 | 8 | 9 | 10 |
| ka1a | 5.854E-01 | 1.018E+00 | 2.608E-02 | 9.137E-01 | 2.505E-01 | 6.583E-01 | 8.256E-02 | 4.916E-01 | 2.804E+00 | 1.209E-01 |
| kd1a | 5.000E-02 | 5.000E-02 | 5.000E-02 | 5.000E-02 | 5.000E-02 | 5.000E-02 | 5.000E-02 | 5.000E-02 | 5.000E-02 | 5.000E-02 |
| ki1 | 5.403E-03 | 2.576E-02 | 1.068E-02 | 1.123E-02 | 5.843E-03 | 7.873E-03 | 7.284E-03 | 1.025E-02 | 2.519E-02 | 3.303E-03 |
| ke1 | 9.657E-05 | 2.356E-04 | 2.576E-04 | 2.524E-05 | 1.353E-05 | 1.599E-05 | 1.788E-05 | 5.427E-06 | 2.994E-05 | 1.522E-05 |
| f_ka1a | 4.250E-02 | 2.148E-02 | 2.071E-02 | 3.878E-02 | 9.525E-02 | 1.620E-02 | 1.156E-02 | 4.217E-01 | 9.999E-01 | 1.340E-02 |
| f_kd1a | 2.625E-02 | 1.000E-02 | 6.686E-01 | 1.396E-02 | 9.097E-02 | 1.793E-02 | 3.747E-02 | 6.843E-01 | 5.618E-02 | 9.981E-01 |
| ka1d | 7.070E-04 | 1.147E-02 | 6.930E-05 | 4.376E-04 | 1.353E-04 | 5.532E-03 | 5.327E-03 | 1.122E-03 | 1.013E-03 | 1.336E-04 |
| ps | 7.951E+03 | 2.368E+03 | 6.744E+05 | 3.681E+05 | 2.878E+04 | 9.531E+03 | 1.154E+04 | 2.670E+05 | 1.932E+01 | 2.816E+05 |
| h3 | 1.000E+00 | 1.000E+00 | 1.000E+00 | 1.000E+00 | 1.000E+00 | 1.000E+00 | 1.000E+00 | 1.000E+00 | 1.000E+00 | 1.000E+00 |
| kdP | 3.162E+00 | 1.123E+00 | 1.079E+00 | 1.691E+00 | 1.599E+00 | 3.162E+00 | 2.968E+00 | 3.160E+00 | 1.011E+00 | 2.797E+00 |
| h2 | 1.500E+00 | 2.000E+00 | 1.641E+00 | 1.955E+00 | 1.502E+00 | 1.685E+00 | 1.998E+00 | 1.867E+00 | 1.501E+00 | 1.500E+00 |
| kdNFKB | 1.034E+00 | 1.308E+00 | 3.161E+00 | 3.096E+00 | 2.304E+00 | 2.773E+00 | 2.087E+00 | 1.434E+00 | 1.639E+00 | 2.174E+00 |
| ka2a | 1.320E-02 | 6.274E-02 | 2.672E-02 | 2.208E-02 | 2.197E-02 | 9.218E+01 | 2.695E-02 | 3.337E-01 | 1.067E-02 | 1.011E-02 |
| kd1d | 6.684E-05 | 6.919E-05 | 1.137E-03 | 6.429E-05 | 7.110E-05 | 2.005E-03 | 8.860E-05 | 8.467E-04 | 6.420E-05 | 6.551E-05 |
| c4a | 5.000E-04 | 5.000E-04 | 5.000E-04 | 5.000E-04 | 5.000E-04 | 5.000E-04 | 5.000E-04 | 5.000E-04 | 5.000E-04 | 5.000E-04 |
| ke2a | 8.353E-02 | 4.194E-02 | 2.341E-02 | 9.911E-02 | 1.361E-02 | 3.751E-03 | 8.649E-03 | 1.177E-02 | 1.549E-02 | 4.211E-03 |
| ki3a | 6.638E-03 | 1.907E-03 | 1.839E-03 | 6.038E-03 | 2.181E-03 | 1.318E-03 | 1.358E-03 | 5.267E-03 | 1.750E-03 | 6.700E-03 |
| ke3a | 2.457E-03 | 1.758E-03 | 7.948E-04 | 9.293E-04 | 1.906E-04 | 1.948E-04 | 6.694E-04 | 3.307E-03 | 3.572E-05 | 9.766E-04 |
| c5a | 2.200E-05 | 2.200E-05 | 2.200E-05 | 2.200E-05 | 2.200E-05 | 2.200E-05 | 2.200E-05 | 2.200E-05 | 2.200E-05 | 2.200E-05 |
| ka | 2.409E-06 | 2.263E-06 | 2.595E-05 | 2.120E-06 | 9.491E-06 | 3.078E-06 | 1.556E-05 | 2.706E-06 | 2.238E-06 | 9.295E-06 |
| ki | 1.156E-02 | 9.935E-03 | 1.130E-02 | 1.129E-02 | 1.138E-02 | 1.157E-02 | 1.160E-02 | 9.715E-03 | 4.903E-03 | 1.160E-02 |
| kp | 6.464E-05 | 7.738E-05 | 2.674E-03 | 9.060E-05 | 2.421E-03 | 1.016E-03 | 1.370E-03 | 6.441E-05 | 1.138E-02 | 8.211E-03 |
| kbA20 | 1.800E-03 | 1.800E-03 | 1.800E-03 | 1.800E-03 | 1.800E-03 | 1.800E-03 | 1.800E-03 | 1.800E-03 | 1.800E-03 | 1.800E-03 |
| c1a | 1.525E+01 | 6.487E+00 | 2.227E+01 | 8.655E+00 | 1.245E+01 | 7.777E+00 | 2.158E+01 | 1.415E+01 | 1.421E+01 | 2.499E+01 |
| h | 2.000E+00 | 2.000E+00 | 2.000E+00 | 2.000E+00 | 2.000E+00 | 2.000E+00 | 2.000E+00 | 2.000E+00 | 2.000E+00 | 2.000E+00 |
| k | 6.681E-02 | 6.515E-02 | 7.179E-02 | 6.586E-02 | 1.263E-01 | 2.033E-01 | 9.796E-02 | 6.501E-02 | 3.657E-01 | 6.579E-02 |
| rs_a | 6.852E-08 | 9.820E-08 | 8.706E-08 | 6.178E-08 | 5.182E-08 | 5.435E-08 | 5.616E-08 | 5.001E-08 | 6.515E-08 | 7.825E-08 |
| c3a | 3.000E-04 | 3.000E-04 | 3.000E-04 | 3.000E-04 | 3.000E-04 | 3.000E-04 | 3.000E-04 | 3.000E-04 | 3.000E-04 | 3.000E-04 |
| c2a | 5.000E-01 | 5.000E-01 | 5.000E-01 | 5.000E-01 | 5.000E-01 | 5.000E-01 | 5.000E-01 | 5.000E-01 | 5.000E-01 | 5.000E-01 |
| k4 | 1.018E-02 | 1.155E-02 | 1.479E-01 | 1.228E-02 | 2.641E-02 | 1.842E-01 | 9.652E-01 | 8.229E-02 | 2.209E-01 | 2.563E-02 |
| c6a | 4.290E-05 | 4.290E-05 | 4.290E-05 | 4.290E-05 | 4.290E-05 | 4.290E-05 | 4.290E-05 | 4.290E-05 | 4.290E-05 | 4.290E-05 |
| c1 | 2.000E-07 | 2.000E-07 | 2.000E-07 | 2.000E-07 | 2.000E-07 | 2.000E-07 | 2.000E-07 | 2.000E-07 | 2.000E-07 | 2.000E-07 |
| k2 | 9.979E-01 | 5.847E-01 | 1.847E-01 | 5.772E-01 | 3.202E-02 | 2.653E-02 | 1.675E-01 | 9.754E-01 | 1.171E-02 | 1.050E-01 |
| c3 | 4.000E-04 | 4.000E-04 | 4.000E-04 | 4.000E-04 | 4.000E-04 | 4.000E-04 | 4.000E-04 | 4.000E-04 | 4.000E-04 | 4.000E-04 |
| c2 | 5.000E-01 | 5.000E-01 | 5.000E-01 | 5.000E-01 | 5.000E-01 | 5.000E-01 | 5.000E-01 | 5.000E-01 | 5.000E-01 | 5.000E-01 |
| c4 | 4.500E-03 | 4.500E-03 | 4.500E-03 | 4.500E-03 | 4.500E-03 | 4.500E-03 | 4.500E-03 | 4.500E-03 | 4.500E-03 | 4.500E-03 |
| kc1a | 4.998E-01 | 3.603E-01 | 6.169E-02 | 4.990E-01 | 1.010E-01 | 2.603E-01 | 1.095E-01 | 3.514E-01 | 8.498E-02 | 2.791E-01 |
| kt1a | 5.796E-03 | 2.681E-05 | 9.794E-05 | 1.160E-06 | 3.056E-03 | 1.410E-06 | 2.146E-06 | 9.194E-04 | 5.631E-03 | 6.875E-06 |
| kc2a | 4.071E-01 | 2.305E-01 | 9.433E-03 | 2.774E-01 | 3.883E-02 | 3.025E-02 | 1.465E-02 | 2.703E-01 | 9.620E-02 | 1.971E-02 |
| kt2a | 1.274E-04 | 1.760E-04 | 3.179E-05 | 1.616E-04 | 5.877E-05 | 7.688E-05 | 2.662E-05 | 2.133E-04 | 3.127E-04 | 3.724E-05 |

**Supplementary Table 4:** Prior boundaries of parameters used for particle swarm optimization of the D2FC and D2FC$^2$ models. D2FC values of N/A represent parameters that were introduced in the D2FC$^2$ model. High and low parameter bounds are equal for parameter values that were held constant during optimization.

| | | | Particle Swarm Parameter Bounds | |
|---|---|---|---|---|
| Name | D2FC Value | Units | Low | High |
| ka1a | 0.5 | 1/(uM*sec) | 1.00E-02 | 1.00E+02 |
| kd1a | 0.05 | 1/sec | 0.05 | 0.05 |
| ki1 | 0.0026 | 1/sec | 0.00026 | 0.026 |
| ke1 | 5.20E-05 | 1/sec | 5.20E-06 | 5.20E-04 |
| f_ka1a | N/A | dimensionless | 0.01 | 1 |
| f_kd1a | N/A | dimensionless | 0.01 | 1 |
| ka1d | N/A | 1/sec | 6.42E-05 | 1.16E-02 |
| ps | N/A | dimensionless | 1 | 1.00E+06 |
| h3 | N/A | dimensionless | 1 | 1 |
| kdP | N/A | uM/sec | 1 | 3.1623 |
| h2 | N/A | dimensionless | 1.5 | 2 |
| kdNFKB | N/A | uM/sec | 1 | 3.1623 |
| ka2a | N/A | 1/( uM *sec) | 1.00E-02 | 1.00E+02 |
| kd1d | N/A | 1/sec | 6.42E-05 | 1.16E-02 |
| c4a | 0.0005 | 1/sec | 0.0005 | 0.0005 |
| ke2a | 0.01 | 1/sec | 1.00E-03 | 1.00E-01 |
| ki3a | 0.00067 | 1/sec | 6.70E-05 | 6.70E-03 |
| ke3a | 0.000335 | 1/sec | 3.35E-05 | 3.35E-03 |
| c5a | 2.20E-05 | 1/sec | 2.20E-05 | 2.20E-05 |
| ka | 2.00E-06 | 1/sec | 1.00E-07 | 1.00E-04 |
| ki | 0.003 | 1/sec | 6.42E-05 | 1.16E-02 |
| kp | 0.0006 | 1/sec | 6.42E-05 | 1.16E-02 |
| kbA20 | 0.0018 | uM | 0.0018 | 0.0018 |
| c1a | 1.00E+00 | dimensionless | 1.00E+00 | 2.50E+01 |
| h | 2 | dimensionless | 2.00E+00 | 2.00E+00 |
| k | 0.065 | uM | 6.50E-02 | 3.00E+00 |
| rs_a | 3.08E-06 | uM/sec | 5.00E-08 | 1.00E-07 |
| c3a | 0.0003 | 1/sec | 0.0003 | 0.0003 |
| c2a | 0.5 | 1/sec | 5.00E-01 | 5.00E-01 |
| k4 | 0.065 | uM | 1.00E-02 | 1 |
| c6a | 4.29E-05 | 1/sec | 4.29E-05 | 4.29E-05 |
| c1 | 2.00E-07 | uM /sec | 2.00E-07 | 2.00E-07 |
| k2 | 0.065 | uM | 1.00E-02 | 1 |
| c3 | 0.0004 | 1/sec | 0.0004 | 0.0004 |
| c2 | 0.5 | 1/sec | 0.5 | 0.5 |
| c4 | 0.0045 | 1/sec | 0.0045 | 0.0045 |
| kc1a | 0.074 | 1/(uM*sec) | 0.009 | 0.5 |
| kt1a | 0.1 | 1/sec | 1.00E-06 | 0.006 |
| kc2a | 0.37 | 1/( uM *sec) | 0.009 | 0.5 |
| kt2a | 0.1 | 1/sec | 1.00E-06 | 0.006 |
