## Supplementary material for "Time-varying stimuli that prolong IKK activation promote nuclear remodeling and mechanistic switching of NF-κB dynamics": D2FCSquared Matlab Files: Helper.m

```
classdef Helper
    properties
        Model 
        SpeciesUnit 
        TimeUnit 
    end 
    
    methods
        %Instantiating the paramters 
        function obj = Helper(Model,SpeciesUnit,TimeUnit)
           obj.Model = Model;
           obj.SpeciesUnit = SpeciesUnit;
           obj.TimeUnit = TimeUnit;
        end

        %Scalar 
        function Scalar(obj,Name,Value,Constant)
            if nargin ==3
                Constant=true;
            end 
            units = "dimensionless";
            addparameter(obj.Model,Name, Value,"ValueUnits",units,"Constant",Constant);
        end 

        function TimeValue(obj,Name,Value,Constant)
            if nargin ==3
                Constant=true;
            end 
            units = obj.TimeUnit;
            addparameter(obj.Model,Name, Value,"ValueUnits",units,"Constant",Constant);
        end 

        %Concentration
        function Concentration(obj,Name,Value)
            if nargin ==3
                Constant=true;
            end 
            units = sprintf("%s",obj.SpeciesUnit);
            addparameter(obj.Model,Name, Value,"ValueUnits",units,"Constant",Constant);
        end 

        %Add a zero order rate constant 
        function ZeroOrder(obj,Name,Value)
            if nargin ==3
                Constant=true;
            end 
            units = sprintf("%s/%s",obj.SpeciesUnit,obj.TimeUnit);
            addparameter(obj.Model,Name, Value,"ValueUnits",units,"Constant",Constant);
        end 

        %Add a first order rate constant 
        function FirstOrder(obj,Name,Value)
            if nargin ==3
                Constant=true;
            end 
            units = sprintf("1/%s",obj.TimeUnit);
            addparameter(obj.Model,Name, Value,"ValueUnits",units,"Constant",Constant);
        end 

        %Add a second order rate constant 
        function SecondOrder(obj,Name,Value)
            if nargin ==3
                Constant=true;
            end 
            units = sprintf("1/(%s*%s)",obj.SpeciesUnit,obj.TimeUnit);
            addparameter(obj.Model,Name, Value,"ValueUnits",units,"Constant",Constant);
        end 
        
        %Add Species
        function AddSpecies(obj,Compartment,Name,InitialValue)
            addspecies(Compartment,Name,InitialValue,"Units",obj.SpeciesUnit);
        end 

        %Add Reaction with mass action kinetics 
        function AddMassActionEquation(obj,Reaction,Parameters)
            %Note: Parameters must be in {} 
            r = addreaction(obj.Model,Reaction);
            addkineticlaw(r,'MassAction','ParameterVariableNames',Parameters);
        end 

        %Adding Reaction with a custom Rate: 
        function AddReaction(obj,Reaction,RateLaw)
            r = addreaction(obj.Model,Reaction);
            r.ReactionRate = RateLaw;

        end 
        
    end 


end 
```
