## Supplementary material for "Time-varying stimuli that prolong IKK activation promote nuclear remodeling and mechanistic switching of NF-κB dynamics": D2FCSquared Matlab Files: D2FCSquared.m

```
function [model,Data] = D2FCSquared()

%==========================================================================
%                        Model Initiation 
%==========================================================================
model = sbiomodel("NFkB Model");


%==========================================================================
%                        Model Compartments
%==========================================================================
Cytoplasm = addcompartment(model,'Cytoplasm',"Capacity",3.33);
Nucleus   = addcompartment(model,'Nucleus',"Capacity",1.0);


helper = Helper(model,"micromole","second");


%==========================================================================
%                Apply Sum of Gaussians for IKK Trajectory 
%==========================================================================
helper.Scalar("a1_coeff",0);
helper.Scalar("b1_coeff",0);
helper.Scalar("c1_coeff",0);
helper.Scalar("a2_coeff",0);
helper.Scalar("b2_coeff",0);
helper.Scalar("c2_coeff",0);
helper.Scalar("a3_coeff",0);
helper.Scalar("b3_coeff",0);
helper.Scalar("c3_coeff",0);
helper.Scalar("a4_coeff",0);
helper.Scalar("b4_coeff",0);
helper.Scalar("c4_coeff",0);


%Sum of Four Guassians for the IKK 
helper.Scalar("IKKSpots",0,false)

sumOfFourGaussians = ['a1_coeff*exp(-((time-b1_coeff)/c1_coeff)^2)' ...
                     '+a2_coeff*exp(-((time-b2_coeff)/c2_coeff)^2)' ...
                     '+a3_coeff*exp(-((time-b3_coeff)/c3_coeff)^2)' ...
                     '+a4_coeff*exp(-((time-b4_coeff)/c4_coeff)^2)'];

addrule(model, "IKKSpots = "+sumOfFourGaussians, 'repeatedAssignment');


%==========================================================================
%                           Model Species
%==========================================================================
helper.AddSpecies(Cytoplasm,"NFkB",0)
helper.AddSpecies(Cytoplasm,"IkBa",0)
helper.AddSpecies(Cytoplasm,"IkBaNFkB",0.3)

helper.AddSpecies(Nucleus,"NFkB",0)
helper.AddSpecies(Nucleus,"NFkBDNA",0)

helper.AddSpecies(Nucleus,"IkBa",0)
helper.AddSpecies(Nucleus,"IkBaNFkB",0)

helper.AddSpecies(Cytoplasm,"pIkBa",0)
helper.AddSpecies(Cytoplasm,"pIkBaNFkB",0)


helper.AddSpecies(Cytoplasm,"IKK",0)
helper.AddSpecies(Cytoplasm,"IKKi",0)
helper.AddSpecies(Cytoplasm,"IKKn",0.8)

helper.AddSpecies(Cytoplasm,"tIkBa",0)
helper.AddSpecies(Cytoplasm,"tCompetitor",0)
helper.AddSpecies(Cytoplasm,"Competitor",0)

helper.AddSpecies(Cytoplasm,"tA20",0)
helper.AddSpecies(Cytoplasm,"A20",0)


%==========================================================================
%                            Parameters
%==========================================================================

parameters_d2fc;


%%%%%%%%%%%% d2fc options %%%%%%%%%%%%%%%%

parameters_d2fc;


helper.Scalar("TR",0,false)

%Kv is the ratio of cytoplasm to nuclear area and needs to stay 
%constant. 
helper.Scalar("kv",kv);


%==========================================================================
%                        Model Reactions
%==========================================================================
%Binding and unbinding of NFkB and IkB in Cytoplasm
helper.SecondOrder('ka1a',ka1a);
helper.FirstOrder("kd1a",kd1a);
helper.AddMassActionEquation( ...
           'Cytoplasm.IkBa + Cytoplasm.NFkB <-> Cytoplasm.IkBaNFkB', ...
            {'ka1a','kd1a'})

%Reaction rates of NFkB transferring between cytoplasm and nucleus while
%accounting for change of volume 
helper.FirstOrder("ki1",ki1);
helper.FirstOrder("ke1",ke1);

[r2,r3,r4,r5]= GenerateCytoplasmNucleurTransfer(model,"NFkB","ki1","ke1","kv");

%Binding and unbinding of NFkB and IkB in Nucleus
helper.Scalar("f_ka1a",0.1)
helper.Scalar("f_kd1a",0.1)

helper.SecondOrder("ka1a_nucleus",1E-14)
helper.FirstOrder("kd1a_nucleus",1E-14)

addrule(model,"ka1a_nucleus = f_ka1a* ka1a","initialAssignment");
addrule(model,"kd1a_nucleus = f_kd1a* kd1a","initialAssignment");
helper.AddMassActionEquation( ...
            'Nucleus.IkBa + Nucleus.NFkB <-> Nucleus.IkBaNFkB', ...
             {'ka1a_nucleus','kd1a_nucleus'})

helper.ZeroOrder("ka1d",1E-3)
helper.Scalar("ps",1)
helper.Scalar("h3",2)
helper.Concentration("kdP",0.5)
NPio = "(1+ps.*((Nucleus.NFkBDNA./kdP).^h3./((Nucleus.NFkBDNA./kdP).^h3 + 1)))"; 


helper.Scalar("h2",1)
helper.Concentration("kdNFKB",0.5)
DCoop = "(Nucleus.NFkB./kdNFKB).^h2./(1+(Nucleus.NFkB./kdNFKB).^h2)";
RateofNFkBBindingDNA = "ka1d*"+DCoop+"*"+ NPio;

helper.AddReaction("Nucleus.NFkB -> Nucleus.NFkBDNA",RateofNFkBBindingDNA)

addobservable(model,"DNACooperativety",DCoop);
addobservable(model,"NPio",NPio);


helper.SecondOrder("ka2a",1E-14)
helper.AddMassActionEquation( ...
            'Nucleus.IkBa + Nucleus.NFkBDNA -> Nucleus.IkBaNFkB', ...
             {'ka2a'});


helper.FirstOrder("kd1d",1E-2)
helper.AddReaction("Nucleus.NFkBDNA ->  Nucleus.NFkB",...
                    "kd1d*Nucleus.NFkBDNA")


%Decay of IkBa in the nucleus and cytplasm
helper.FirstOrder("c4a",c4a)
helper.AddMassActionEquation("Nucleus.IkBa -> null",{'c4a'})
helper.AddMassActionEquation("Cytoplasm.IkBa -> null",{'c4a'})

%Exit of NFkB/IkB complex from the nucelus to the Cytoplasm 
helper.FirstOrder("ke2a",ke2a);
r7 = RateExitNucleus(model,"IkBaNFkB","ke2a","kv");

%Reaction rates for IkB transferring between the cytoplasm and nucleus
%while accounting for change of volume 
helper.FirstOrder("ki3a",ki3a);
helper.FirstOrder("ke3a",ke3a);
[r8,r9,r10,r11] = GenerateCytoplasmNucleurTransfer(model,"IkBa","ki3a","ke3a","kv");

%IkB Cytoplasmic Degragation on NFkB. NFkB is generated 
helper.FirstOrder("c5a",c5a);
helper.AddMassActionEquation( ...
                             "Cytoplasm.IkBaNFkB -> Cytoplasm.NFkB", ...
                             {'c5a'})

%Neutral IKK (IKKn) converting to active IKK (IKK) 
helper.FirstOrder("ka",1E-3);
helper.AddReaction( ...
         "Cytoplasm.IKKn -> Cytoplasm.IKK",...
         "TR*IKKSpots*ka*Cytoplasm.IKKn")

%IKK converting to inactive IKKn
helper.FirstOrder("ki",ki)
helper.AddMassActionEquation("Cytoplasm.IKK -> Cytoplasm.IKKi",{'ki'})

%Inactive IKK (IKKn) converting to neutural IKK (IKKn)
helper.FirstOrder("kp",kp)
helper.Concentration("kbA20",kbA20)
helper.AddReaction("Cytoplasm.IKKi -> Cytoplasm.IKKn", ...
    "kp*Cytoplasm.IKKi*kbA20/(kbA20+A20)")


%Formation of IkBa mRNA transcript in response to NFkB in the nucleus 
helper.Scalar("c1a",0.5);
helper.Scalar("h",h);
helper.Concentration("k",k);
helper.ZeroOrder("rs_a",3.0800e-06)
RateOfIkBaTranscription = "rs_a.*(1 + c1a.*(Nucleus.NFkB./k).^h ./( (Nucleus.NFkB./k).^h + 1))";
helper.AddReaction("null -> Cytoplasm.tIkBa",RateOfIkBaTranscription)

%Decay of IkBa mRNA transcript. Follows basic mass action kinetics 
helper.FirstOrder("c3a",c3a);
helper.AddMassActionEquation("Cytoplasm.tIkBa -> null",{'c3a'})

%Transcript of IkBa can make IkBa in the cytoplasm 
helper.FirstOrder("c2a",c2a)
helper.AddReaction("null -> Cytoplasm.IkBa",...
                   "c2a*Cytoplasm.tIkBa")


%Competitor Transcript Formation 
helper.Concentration("k4",k4)
helper.AddReaction("null -> Cytoplasm.tCompetitor",...
 "rs_a* c1a* (Nucleus.NFkB/k)^(h+1) /( (Nucleus.NFkB/k)^(h+1) + (Competitor/k4)^(h+1)  + 1)")

%Competitor Transcript Decay 
helper.FirstOrder("c6a",c6a)
helper.AddMassActionEquation("Cytoplasm.tCompetitor -> null",{'c6a'})

%Competitor Protein Creation 
helper.AddReaction("null -> Cytoplasm.Competitor", ... 
              "c2a*Cytoplasm.tCompetitor")

%Competitor Protein Decay 
helper.AddMassActionEquation("Cytoplasm.Competitor -> null",{'c4a'})

%A20 transcript formation [tA20]
helper.ZeroOrder("c1",c1)
helper.Concentration("k2",k2)
helper.AddReaction("null -> Cytoplasm.tA20",...
 "c1* (Nucleus.NFkB/k)^(h+1) /( (Nucleus.NFkB/k)^(h+1) + (Cytoplasm.Competitor/k2)^(h+1)  + 1)")

%A20 Trasncript Decay 
helper.FirstOrder("c3",c3)
helper.AddMassActionEquation("Cytoplasm.tA20 -> null",{'c3'})

%A20 Creation 
helper.FirstOrder("c2",c2)
helper.AddReaction("null -> Cytoplasm.A20", ... 
              "c2*Cytoplasm.tA20")

%A20 Protein Decay 
helper.FirstOrder("c4",c4)
helper.AddMassActionEquation("Cytoplasm.A20 -> null",{'c4'})

%Phosphorlating IkBa 
helper.SecondOrder("kc1a",kc1a)
helper.AddReaction("Cytoplasm.IkBa -> Cytoplasm.pIkBa", ...
                  "kc1a*Cytoplasm.IKK*Cytoplasm.IkBa")

%Decay of pIkBa 
helper.FirstOrder("kt1a",kt1a)
helper.AddMassActionEquation("Cytoplasm.pIkBa -> null",{'kt1a'})

%Phosphorlating pIkBaNFkB 
helper.SecondOrder("kc2a",kc2a)
helper.AddReaction("Cytoplasm.IkBaNFkB -> Cytoplasm.pIkBaNFkB", ...
                  "kc2a*Cytoplasm.IKK*Cytoplasm.IkBaNFkB")

%Phosphorlated pIkBaNFkB will break apart where IkBa will decay and NFkB
%will be released 
helper.FirstOrder("kt2a",kt2a)
helper.AddMassActionEquation("Cytoplasm.pIkBaNFkB -> Cytoplasm.NFkB", ...
                            {'kt2a'})


%==========================================================================
%Observables for model output 
%==========================================================================
addobservable(model,"TotalNFkB_Cyt","Cytoplasm.NFkB + Cytoplasm.IkBaNFkB + Cytoplasm.pIkBaNFkB");
addobservable(model,"TotalNFkB_Nuc","Nucleus.NFkB +Nucleus.IkBaNFkB+ Nucleus.NFkBDNA ");
addobservable(model,"FractionNuc2Cyt","(TotalNFkB_Nuc./kv)./TotalNFkB_Cyt");
addobservable(model,"RelativeNFkB_Nuc","TotalNFkB_Nuc./TotalNFkB_Nuc(1)");

addobservable(model,"Total_IkBalphaCytoplasm","Cytoplasm.IkBa + Cytoplasm.IkBaNFkB");
addobservable(model,"Total_IkBalphaPhoshorlated","Cytoplasm.pIkBa + Cytoplasm.pIkBaNFkB");


addobservable(model,"Total_IkBalphaNucleus","Nucleus.IkBa + Nucleus.IkBaNFkB");
addobservable(model,"FractionIkBaCytoplasm","Total_IkBalphaCytoplasm./(Total_IkBalphaCytoplasm) ");
addobservable(model,"FractionIkBaNucleus","Total_IkBalphaNucleus./(Total_IkBalphaNucleus) ");


%Need to Log Parameter States 
configsetObj = getconfigset(model);
% Get the current StatesToLog
currentStatesToLog = configsetObj.RuntimeOptions.StatesToLog;

pars2Log = {'ps','kdP', 'h3','c1a','k','h','kv','NFkBStability','nUnbinding','kdNFKB','h2'};
for i = 1:length(pars2Log)
    ithParameter = pars2Log{i};
    ithParameter = sbioselect(model,"Type","parameter","Name",ithParameter);
    currentStatesToLog = [currentStatesToLog; ithParameter];
end 
 configsetObj.RuntimeOptions.StatesToLog = currentStatesToLog;

end 
```
