## Supplementary material for "Time-varying stimuli that prolong IKK activation promote nuclear remodeling and mechanistic switching of NF-κB dynamics": D2FCSquared Matlab Files: GenerateCytoplasmNucleurTransfer.m

```
function [Rate1,Rate2,Rate3,Rate4] = GenerateCytoplasmNucleurTransfer(Model,SpeciesName,KEntry,KExit,NuclearScalingFactor)
    [Rate1,Rate2] = RateIntoNucleus(Model,SpeciesName,KEntry,NuclearScalingFactor);
    [Rate3,Rate4] = RateExitNucleus(Model,SpeciesName,KExit,NuclearScalingFactor);
   
end 
```
