## Supplementary material for "Time-varying stimuli that prolong IKK activation promote nuclear remodeling and mechanistic switching of NF-κB dynamics": D2FCSquared Matlab Files: GetMeanNuclearRelAExpData.m

```
function [meanRelAFC] = GetMeanNuclearRelAExpData(Data,ScenarioName)
    scenarios = ["Control" "1X30 sec" "1X2 min" "1X6 min" "1X15 min"...
             "1X30 min" "2X3 min" "3X2 min" "4X1.5 min"]; 

    keyValue = ["IKK_RelA_negativeControl"
                "IKK_RelA_Single_30sec_pulse"
                "IKK_RelA_Single_2min_pulse"
                "IKK_RelA_Single_6min_pulse"
                "IKK_RelA_Single_15min_pulse"
                "IKK_RelA_Single_30min_pulse"
                "IKK_RelA_Two_3min_pulses_05min_gap"
                "IKK_RelA_Three_2min_pulses_05min_gap"
                "IKK_RelA_Four_90sec_pulses_05min_gap"]; 

    idx = find(ScenarioName ==scenarios ); 

    keyScenario = keyValue(idx); 

    relAFC = Data.(keyScenario).RelA_FC;
    
    meanRelAFC = mean(relAFC,2);

end 
```
