## Supplementary material for "Time-varying stimuli that prolong IKK activation promote nuclear remodeling and mechanistic switching of NF-κB dynamics": D2FCSquared Matlab Files: RateExitNucleus.m

```
function [RateExit,RateEnter] = RateExitNucleus(Model,SpeciesName,K,ScalingFactor)
    reaction1 = sprintf("null -> Cytoplasm.%s" ,SpeciesName);
    reactionRate1 =  sprintf("%s * Nucleus.%s",K,SpeciesName);

    reaction2 = sprintf("Nucleus.%s -> null"   ,SpeciesName);
    reactionRate2 =  sprintf("%s * %s * Nucleus.%s",ScalingFactor,K,SpeciesName);


    
    RateEnter = addreaction(Model,reaction1,"ReactionRate", reactionRate1);
    RateExit = addreaction(Model,reaction2 ,"ReactionRate",reactionRate2);

end
```
