## Supplementary material for "Time-varying stimuli that prolong IKK activation promote nuclear remodeling and mechanistic switching of NF-κB dynamics": D2FCSquared Matlab Files: RateIntoNucleus.m

```
function [RateExit,RateEnter] = RateIntoNucleus(Model,SpeciesName,K,ScalingFactor)
    reaction1 = sprintf("Cytoplasm.%s -> null"   ,SpeciesName);
    reaction2 = sprintf("null -> Nucleus.%s"   ,SpeciesName);


    reactionRate1 = sprintf("%s * Cytoplasm.%s",K,SpeciesName);
    reactionRate2 = sprintf("%s * %s * Cytoplasm.%s",ScalingFactor,K,SpeciesName);

    RateExit = addreaction(Model,reaction1,"ReactionRate", reactionRate1);
    RateEnter = addreaction(Model, reaction2,"ReactionRate", reactionRate2);
end
```
