## Supplementary material for "Time-varying stimuli that prolong IKK activation promote nuclear remodeling and mechanistic switching of NF-κB dynamics": D2FCSquared Matlab Files: SimulateModel.m

```
function [simdata,names,t] = SimulateModel(model,IntialCondition)


%==========================================================================
%                        Simulate the Model 
%==========================================================================

csObj = getconfigset(model,'active');
% Change solver type
set(csObj, 'SolverType', 'sundials'); 

% Change tolerances
set(csObj.SolverOptions, 'AbsoluteTolerance', 1e-8); % Change absolute tolerance
set(csObj.SolverOptions, 'RelativeTolerance', 1e-8); % Change relative tolerance

duration = 10; %Days 
durationSeconds = duration*24*60*60;
set(csObj,'Stoptime',durationSeconds);


%I need to make sure the inital values always stay the same. I found some
%evidence that they may be changing during fit due to failing simulation
%and not resetting the intial values. 
for i = 1:numel(model.Species)
    model.Species(i).InitialAmount = IntialCondition(i);
end

[t, simdata, names] = sbiosimulate(model);

for i = 1:numel(model.Species)
    newInitial = simdata(end,i);

    %The if statement prevents setting inital values that are slightly less
    %than zero due to numerical error. 
    if newInitial<1E-14
        newInitial = 0;
    end 

    model.Species(i).InitialAmount = newInitial;
end


%==========================================================================
%                         Simulate Simulated Cells  
%==========================================================================

TR = sbioselect(model,"Type","parameter",'Name',"TR");

TR.value = 1; %Turning on Stimulation 

stopTimeHours = 3;
set(csObj,'Stoptime',stopTimeHours*60*60);

timeVector = [0:4*60:stopTimeHours*60*60];
set(csObj.SolverOptions, 'MaxStep', 2*60);
[t, simdata, names] = sbiosimulate(model);

%resetIntialValues
for i = 1:numel(model.Species)
    model.Species(i).InitialAmount = IntialCondition(i);
end

TR.value = 0; %Turning Off Stimulation
end 
```
