## Supplementary material for "Time-varying stimuli that prolong IKK activation promote nuclear remodeling and mechanistic switching of NF-κB dynamics": D2FCSquared Matlab Files: UpdateParameters.m

```
function [model] = UpdateParameters(model,parameters2Update,StartIndex)

%Updating Model parameters starting from an index. 
lengthOfParameters = length(parameters2Update);
for i = 1:lengthOfParameters
    index2Update = i+StartIndex-1;
    model.Parameters(index2Update).value = parameters2Update(i);
end 

end 
```
