## Supplementary material for "Time-varying stimuli that prolong IKK activation promote nuclear remodeling and mechanistic switching of NF-κB dynamics": D2FCSquared Matlab Files: parameters_d2fc.m

```
%---------------------------------------%
% Parameter values & initial conditions %
%---------------------------------------%

% ----------------------------------------------------------------------- %
% ------------------------ WT or A20 KO cell type ----------------------- %
A20On=1;            % A20 (=1) or A20 knockouts (=0)

% ------------------------------- Volume -------------------------------- %
% CHECK SCALING OF ALL PARAMETERS IF SCANNING KV, e.g. on transcription 
kv = 3.3;                   % NEW - experimentally determined
tv = 2700;                  % NEW - experimentally determined
nScale = kv+1;              % Convert from total cell to nuclear volume
cScale = (1/kv)+1;          % Convert from total cell to cytoplasmic volume

% -------------- Scaling constants for mol/conc conversion -------------- %
avogadro = 6.022*10^23;
cellMolecules = avogadro*(tv*10^-15)*10^-6;     % 1uM concentration as molecules/cell
cScaleMol = cellMolecules/cScale;   % to convert to compartmental numbers
nScaleMol = cellMolecules/nScale;   % to convert to compartmental numbers

% ---------------- Initial conditions (uM & molecules) ------------------ %
% For ~100,000 molecules in cytoplasmic compartment
% (100000/cellMolecules)*cScale = 0.0801 uM

ye=zeros(1,17);             % All species initialised to zero
NF = 0.08;                  % Total cell NFkB as cytoplasmic concentration
ye(3) =NF;                  % NF-kB is initialised to the cytoplasm bound to IkBa

% ----------------------------------------------------------------------- %
% ------------------ ODE Parameters (s-1, uM-1 s-1)---------------------- %

% IKK %
ye(8) = 0.08;               % Initialise total IKK (uM)
kp = 0.0006;                % IKKn production rate
ka = 0.000002;              % Activation caused by TNF; **for D2FC, ka is assigned with the "Doses" array
ki = 0.003;                 % Spontanouous IKK inactivation


% A20 protein %
c4 = 0.0045;                 % A20 degradation rate
kbA20 = 0.0018;              % Half-max A20 inhibition concentration (uM)

% IKK-IkBa-NFkB interactions %
ka1a = 0.5;                  % NFkB-IkBa association (Hoffmann et al. 2002)
kd1a = 0.05;                 % NFkB-IkBa dissociation *In D2FC, this is adjusted from the value described previously (Hoffmann et al. 2002)


% IKK phosphorylation of IkBa and IkBaNFkB 
IKKafold = 2;                % IkBa degradation
kc1a = 0.037*1*IKKafold;     % catalysis of IKK.IkBa dimer
kc2a = 0.037*5*IKKafold;     % catalysis of IKK.IkBa.NFkB trimer

% Signal/IKK dependent IkBa degradation %
kt1a = 0.1;                  % Degradation of IkBa (IKK dependent from dimer)
kt2a = 0.1;                  % Degradation of IkBa (IKK dependent from trimer)


% Constitutive IkBa degradation %
c4a  = 0.0005;            % 0.0005 'apparent' free cytoplasmic/nuclear IkBa degradation (t1/2 60-110 min) (Rice & Ernst, 1993; Pando & Verma, 2000)
c5a  = 0.000022;          % NFkB-Complexed cytoplasmic/nuclear IkBa degradation (t1/2 530-550 min) (Pando & Verma, 1993)

% Transport %
ki1  = 0.0026;            % Observed SK-N-AS cell range: avg (0.0026 � 0.0018 s-1 NFkB nuclear import
ke1 = ki1/50;             % NFkB nuclear export (derived from experimentally determined NFkB import - import/50 (Carlotti et al., 2000))
ke2a = 0.01;              %  Free: NFkB.IkB complex nuclear export - Fitted
ki3a  = 0.00067;          % Observed SK-N-AS cel range: (avg - max): 0.00043 � 0.00024 s-1 IkBa nuclear import
ke3a  = ki3a/2;           % IkBa nuclear export (derived from experimentally determined IkBa import - import/2 (Carlotti et al., 2000))

% NFkB-dependent synthesis %
h = 2;                    % 2 Order of hill function controlling transcription *for tA20, tComp and tInducedTarget, the order of the hill function is h+1 (i.e. h+1 = 3); see nfkbmodel.m 
k = 0.065;                % for h=1, k=0.043, h=2, k=0.065  Constant representing nNFkB concentration at which half-maximal transcription rate occurs (uM)
k2 = 1*k;                 % Activation coefficient for competitor on "A20" promoter(The concentration of competitor protein required to significantly [occupy promoter]or [repress expression] of target)
k3 = 0.5*k;               % Activation coefficient for competitor on "InducedTarget" promoter(The concentration of competitor protein required to significantly [occupy promoter]or [repress expression] of target)[scanned parameter] 
k4 = 1*k;                 % Activation coefficient for competitor on its own promoter(The concentration of competitor protein required to significantly [occupy promoter]or [repress expression] of target)[scanned parameter] 

% IkBa synthesis %
c1a = (1.4*10^-7);        % Inducible IkBa mRNA synthesis
c2a  = 0.5;               % IkBa translation rate (Lipniacki et al., 2004)
c3a = 0.0003;             % IkBa mRNA degradation

% A20 synthesis %
c1 = ((2*10^-7)*A20On);   % Inducible A20 mRNA synthesis
c2 = 0.5;                 % A20 mRNA translation rate (Lipniacki et al., 2004)
c3 = 0.0004;              % 0.00048 original A20 mRNA degradation
%Competition
Comp = 1;                 %
c6a = c3a/7;              % tCompetitor degradation

TR = 0;
% ----------------------------------------------------------------------- %
% ------------ For Output and function passing/remote updating ---------- %

% --- parameter array
p_d2fc=[kv,tv,kp,ka,ki,ka1a,kd1a,kc1a,kc2a,kt1a,kt2a,c4a,c5a,ki1,ke1,ke2a,ki3a,ke3a,... % 1-18
   h,k,c1a,c2a,c3a,c1,c2,c3,c4,kbA20,k2,k3,k4,Comp, c6a, TR]; % 19 - 33

% --- String array of parameters for plotting
parnames=["kv     ";"tv     ";"kp     ";"ka     ";"ki     ";"ka1a   ";"kd1a   ";"kc1a   ";"kc2a   ";"kt1a   ";"kt2a   ";"c4a    ";"c5a    ";"ki1    ";"ke1    ";"ke2a   ";"ki3a   ";"ke3a   "; ... % 1 - 18
"h      ";"k      ";"c1a    ";"c2a    ";"c3a    ";"c1     ";"c2     ";"c3     ";"c4     ";"kbA20  ";"k3  "];  % 19 - 29
```
