## Supplementary material for "Time-varying stimuli that prolong IKK activation promote nuclear remodeling and mechanistic switching of NF-κB dynamics": D2FCSquared Matlab Files: RunAverageIKKTrajectory.m

```
clc;clear;

addpath("Classes/")
addpath("Data/")
addpath("Functions/")

%==========================================================================
%                        User Input
%==========================================================================
%Note: The file will run with the following warning "Warning: Reported from
%Dimensional Analysis: This is expected behavior from the simulation and 
% the model is running correctly."

%Define Model to Choose: 
%   1 = D2FC Squared
%   2 = D2FC Optimized 
%   3 = D2FC (Orginal Parameters) 
%   4 = Custom (Default D2FCSquared) 
modelType = 1;  


%==========================================================================
%                        Model Loading and Simulation 
%==========================================================================
fittedIKKProfiles = readtable("Data/MeanIKKTrajectories.csv");
scenarios = fittedIKKProfiles.Scenarios; 
parameters = readtable("ModelParameters.xlsx");

switch modelType
    case 1
        disp("D2FC Squared Model Applied")
        displayTitle = "D2FC^2 Model";
        parameterSet = parameters.D2FCSquared;
    case 2
        disp("D2FC Optimized Model Applied")
        displayTitle = "D2FC Optimized Model";
        parameterSet = parameters.D2FCOptimized;
    case 3
        disp("D2FC Model Applied")
        displayTitle = "D2FC Model";
        parameterSet = parameters.D2FC;
    case 4
        disp("Custom Model Applied")
        displayTitle = "Customized Model";
        parameterSet = parameters.Custom;
    otherwise
        error("Invalid model type. Please choose a value between 1 and 4.");
end


model = D2FCSquared(); 
model = UpdateParameters(model,parameterSet,16);
intialCondition = [model.Species.Value];


%% ==========================================================================
%                       Simulating Mean IKK Trajectories 
%==========================================================================

load("Data/ExpData.mat");

figure 
hold on 
colors = [
    0, 0.4470, 0.7410;    % Blue
    0.8500, 0.3250, 0.0980; % Red
    0.9290, 0.6940, 0.1250; % Yellow
    0.4940, 0.1840, 0.5560; % Purple
    0.4660, 0.6740, 0.1880; % Green
    0.3010, 0.7450, 0.9330; % Cyan
    0.6350, 0.0780, 0.1840; % Magenta
    0.8500, 0.3250, 0.0980; % Orange
    0.0000, 0.4470, 0.7410; % Dark Blue
];

for ith = 1:4
    ithScenarios = scenarios{ith}; 
    expData = GetMeanNuclearRelAExpData(ExpData,ithScenarios);
    ithIKKPars = fittedIKKProfiles{ith,2:end};
    model = UpdateParameters(model,ithIKKPars,1);
    [simdata,names,t] = SimulateModel(model,intialCondition); 
    idxFoldChange = find("RelativeNFkB_Nuc" == names);      
    nuclearRelAFC = simdata(:,idxFoldChange);
    plot(t./60,nuclearRelAFC,DisplayName=ithScenarios,Color=colors(ith,:))
    scatter(0:4:181,expData,MarkerEdgeColor=colors(ith,:),DisplayName="Data:"+ithScenarios)
    legend()
    xlabel("Time [Minutes]","FontSize",20)
    ylabel("Nuclear RelA [Fold Change]","FontSize",20)
    title("Fitting: "+displayTitle,FontSize=20)
    ylim([0 5])
end 


figure 
hold on 
for ith = 5:9
    ithScenarios = scenarios{ith}; 
    ithIKKPars = fittedIKKProfiles{ith,2:end};
    expData = GetMeanNuclearRelAExpData(ExpData,ithScenarios);
    model = UpdateParameters(model,ithIKKPars,1);
    [simdata,names,t] = SimulateModel(model,intialCondition); 
    idxFoldChange = find("RelativeNFkB_Nuc" == names);      
    nuclearRelAFC = simdata(:,idxFoldChange);
    plot(t./60,nuclearRelAFC,DisplayName=ithScenarios,Color=colors(ith,:))
    scatter(0:4:181,expData,MarkerEdgeColor=colors(ith,:),DisplayName="Data:"+ithScenarios)
    legend()
    xlabel("Time [Minutes]","FontSize",20)
    ylabel("Nuclear RelA [Fold Change]","FontSize",20)
    title("Validation: "+displayTitle,FontSize=20)
    ylim([0 5])
end 
```
