## Supplementary material for "Time-varying stimuli that prolong IKK activation promote nuclear remodeling and mechanistic switching of NF-κB dynamics": D2FCSquared Matlab Files: RunSingleCellTrajectories.m

```
clc;clear;

addpath("Classes/")
addpath("Data/")
addpath("Functions/")

%==========================================================================
%                        User Input
%==========================================================================
%Note: The file will run with the following warning "Warning: Reported from
%Dimensional Analysis: This is expected behavior from the simulation and 
% the model is running correctly."

%Define Model to Choose: 
%   1 = D2FC Squared
%   2 = D2FC Optimized 
%   3 = D2FC (Orginal Parameters) 
%   4 = Custom (Default D2FCSquared) 
modelType = 1;  


%==========================================================================
%                        Model Loading and Simulation 
%==========================================================================
fittedIKKProfiles = readtable("Data/MeanIKKTrajectories.csv");

parameters = readtable("ModelParameters.xlsx");

switch modelType
    case 1
        disp("D2FC Squared Model Applied")
        displayTitle = "D2FC^2 Model";
        parameterSet = parameters.D2FCSquared;
    case 2
        disp("D2FC Optimized Model Applied")
        displayTitle = "D2FC Optimized Model";
        parameterSet = parameters.D2FCOptimized;
    case 3
        disp("D2FC Model Applied")
        displayTitle = "D2FC Model";
        parameterSet = parameters.D2FC;
    case 4
        disp("Custom Model Applied")
        displayTitle = "Customized Model";
        parameterSet = parameters.Custom;
    otherwise
        error("Invalid model type. Please choose a value between 1 and 4.");
end


model = D2FCSquared(); 
model = UpdateParameters(model,parameterSet,16);
intialCondition = [model.Species.Value];

%==========================================================================
%                 Loading Single Cell trajectories 
%==========================================================================

locOfParmeterSet = "Data/SingleCellTrajectories.xlsx"; 

scenarios = ["Control" "1X30 sec" "1X2 min" "1X6 min" "1X15 min"...
             "1X30 min" "2X3 min" "3X2 min" "4X1.5 min"]; 

ikkParameterFits = {}; 


for ith = 1:length(scenarios)
    ithScenario = scenarios{ith}; 
    ithSingleCellFits = readtable(locOfParmeterSet,"Sheet",ithScenario);
    ikkParameterFits{end+1} = table2array(ithSingleCellFits); 
end 

%% ========================================================================
%                 Simulate Single Cell Scenarios 
%==========================================================================
simulationData = []; 
expTimes = 0:4:181;
nTimePoints = length(expTimes); 

nuclearNFkBFoldChange = {}; 
for ith = 1:length(scenarios)
    ithScenario = scenarios{ith};
    disp("Simulating "+ithScenario)
    ikkPars = ikkParameterFits{ith}; 
    nCells = size(ikkPars,1);

    ithFoldChangeStore = zeros(nTimePoints,nCells);
    for jth = 1:nCells
        jthIKKTraj = ikkPars(jth,:);
        model = UpdateParameters(model,jthIKKTraj,1);
        [simdata,names,t] = SimulateModel(model,intialCondition); 
        idxFoldChange = find("RelativeNFkB_Nuc" == names);
        
        nuclearRelAFC = simdata(:,idxFoldChange);
        ithFoldChangeStore(:,jth) = interp1(t./60, nuclearRelAFC, expTimes, 'linear'); 
    end 
    nuclearNFkBFoldChange{end+1}= ithFoldChangeStore;
end 


%% ========================================================================
%                 Plotting Single cell Trajectories
%==========================================================================

figure 
set(gcf, 'Position', [100, 100, 1900, 300]);  % Set figure to be square
t = tiledlayout(1,length(scenarios));
for ith = 1:length(scenarios)
    ithScenario = scenarios{ith};
    nexttile()
    plot(expTimes,nuclearNFkBFoldChange{ith},"Color",[0 0 0 0.1])
    ylim([0 6])
    if ith == 1
        ylabel("Nuclear RelA (Fold Change)","FontSize",20)
    else 
        yticklabels([]);
    end 
    xlabel("Time [Minutes]","FontSize",15)
    title(ithScenario,FontSize=20)
end 
t.Padding="compact"; 
t.TileSpacing = "compact";
sgtitle(displayTitle,fontsize=30);
```
