## Supplementary material for "Time-varying stimuli that prolong IKK activation promote nuclear remodeling and mechanistic switching of NF-κB dynamics": D2FCSquared Matlab Files: ReadMe.docx

The contents contain the Matlab files needed to run the D2FC^2^, D2FC optimized, and D2FC models, as described in the main paper. Files were run using Matlab 2023a with SimBiology (Version 6.4.1).RunAverageIKKTrajectory m-file will simulate the nuclear Rela (fold change) trajectories for the average data used for fitting and validation. RunSingleCellTrajectories can simulate all the individual single-cell trajectories and output a graph of the results. ModelParameters.xlsx is an excel file containing the parameter sets for each model. The last column labeled ‘Custom’ is intended to be a place where the user can modify parameters individually where the default parameters are the D2FC^2^. The file parameters_d2fc.m contains the original parameter values and sources used in the D2FC model. The D2FCSquared.m contains the mechanistic model using command line tools in SimBiology to create the model. The D2FCSquared model can be converted to D2FC by setting the appropriate parameters to 0.

The Data folder contains three items, a ExpData.mat folder containing the experimental time course trajectories of nuclear RelA (fold change) and IKK spots for all experimental conditions. The MeanIKKTrajectories.csv file contains the parameters that best fit a sum of four Gaussian equations to the mean experimental NEMO spot trajectories. SingleCellTrajectories.xlsx is an Excel sheet that includes the best-fit parameters for each NEMO spot trajectory. Each tab represents an experimental condition. These parameters are used to run a simulation for each condition.

The Helper.m file is a custom class developed to aid in the model input. Using SimBiology command line tools leads to repetition and the opportunity for user input errors, where the Helper class creates a less error prone wrapper around the SimBiology command line tools. In the functions folder includes files for both generating the model and simulating. SimulateModel simulates the model to steady state without NEMO spot formation for 10 simulated days, followed by 3 hours with NEMO spot formation, consistent with experimental observations.

GetMeanNuclearRelAExpData, RateExitNucleus, and RateIntoNucleus are functions to help with the creation of the ODEs for the transfer of analytes from the cytoplasm and nucleus. These functions were created to ensure consistency across all species that needed the same set of equations. UpdateParameters is a function that can update the model parameters from an ordered list of values (see ModelParameters.xlsx). GetMeanNuclearRelAExpData is a function that extracts experimental data from the variable format it is in.
